# Asymmetric Reinforcement Learning Explains Human Choice Patterns in Decision-making Under Risk

**DOI:** 10.64898/2026.03.09.710615

**Authors:** Niloufar Shahdoust, Rhiannon L. Cowan, T. Alexander Price, Tyler S. Davis, Andrew Liu, Rikki Rabinovich, Veronica Zarr, Mark Libowitz, Ben Shofty, Shervin Rahimpour, Alla Borisyuk, Elliot H. Smith

## Abstract

Human decisions under uncertainty are shaped by experience, but the computations that translate expectation and experience into choice remain debated in neural and cognitive science. Prior studies highlight reinforcement learning (RL) as a unifying framework, yet it is unclear whether human behavior under risk is better captured by symmetric updating from outcomes or by asymmetric learning that weights reward and loss differently. This work examines which learning strategies better explain trial-by-trial choices given contextual uncertainty and manipulations of outcome distributions. Our results show that a Risk Sensitive (RS) model with asymmetric learning rates best explains human behavior in our novel decision-making task. Fitting candidate models to individual trial histories yielded value signals that predicted both choice and response time. These results highlight that RS model, as an asymmetric learning provides a concise and identifiable account of behavior in decision-making under risk tasks.

## 1 Introduction

Humans constantly predict potential rewards and losses when making decisions with uncertain outcomes [1]. Yet the computational mechanisms of this process, especially how learning from positive versus negative outcomes shapes decision-making under risk, remain poorly understood. Individuals often arbitrate among learning strategies in different situations, often giving more weight to whichever strategy has been more reliable in recent trials, and they differ substantially in which strategies they tend to rely on most [2]. As contextual uncertainty increases, models that weight strategies by uncertainty, shift in behavior from value-based choice toward frequency-based responding [3], and when evidence is asymmetric or rare, people adopt diverse strategies that can even invert the classic bias–variance trade-off [4]. More broadly, statistical regularities in the environment can bias reward learning by shifting attention and modulating learning rates [5]. Consistent patterns appear in deep RL [6], where heterogeneous navigation strategies are linked to learned representations [7].

Across development, experience-based learning under risk varies systematically, with adolescents learning less efficiently, making more stochastic choices, and displaying heterogeneous strategies relative to adults [8]. Previous studies found that reward prediction error (RPE)-based learning is not restricted to the prediction of reward, but also to risk [9–11]. Beyond guiding choice strategies, RPE also shapes learning, an observation that situates RL variables as candidate signals for future neural analyses [12, 13]. Neuroimaging further shows that striatal RPE signals are themselves risk-sensitive during learning [14]. In experience-based risky choice, differential learning from reward versus loss, together with nonlinear utility, predicts risk preferences, motivating a focus on asymmetric learning [15]. Together, these findings motivate testing which learning rules better explain human trial-level behavior under varying risk and uncertainty.

Heterogeneity in decision strategies across individuals, makes it difficult to deter-mine which candidate computational model best explains their behavior [2]. Classic work on decisions from experience shows that rare events are systematically under-weighted, challenging symmetric updating assumptions and raising questions about how value is learned in risky environments [16, 17]. Differences in learning from rewards versus losses are often difficult to interpret because their behavioral signatures can arise from multiple underlying mechanisms, making careful model comparison essential [18, 19]. Although prior work has proposed risk-sensitive learning signals, the behavioral learning rule that best links trial-wise value differences to both choice behavior and response time (RT) remains unresolved [14, 20]. Together, these challenges highlight the need for interpretable, trial-level learning variables that robustly capture individual differences in behavior across risky decision contexts. To address this gap, candidate learning models were compared to identify the model whose latent variables best predict trial-by-trial choice and RT in experience-based risky decision-making.

In this work, five candidate RL models were fit to participants’ trial-by-trial choices in a novel static risky decision-making task: (1) Win-Stay/Lose-Shift (WSLS) as a simple heuristic model, (2) Rescorla–Wagner [21] with an ϵ-Greedy policy [6] emphasizing exploitation with occasional exploration, (3) Rescorla–Wagner with a Softmax [6, 22] policy that balances exploration and exploitation, (4) a Risk-Sensitive (RS) model with asymmetric learning rates and (5) a Dual-Q model with separate Q-value updates for explicit reward and risk. Models (1)–(3) capture symmetric learning from rewards and losses while using different policies, whereas model (4) explicitly tests asymmetric learning components in human behavior, and model (5) incorporates an explicit parameterization of risk alongside the reward, albeit separately. Candidate models for experience-based risky decision-making were compared to identify interpretable models that capture trial-level variation in choice and RT across participants.

Understanding how individuals’ learning and decision-making strategies have direct relevance in mental health and addiction research. Disorders such as gambling disorder and substance use disorder (SUD) are characterized by altered sensitivity to reward and punishment, contributing to compulsive and risky behavior [23, 24]. Computational psychiatry frameworks argue that formal RL models provide mechanistic insight into psychiatric dysfunction by quantifying latent variables such as RPE signaling and value updating [25, 26]. In gambling disorder, impaired integration of negative feedback and altered value learning have been documented, suggesting asymmetric learning from gains and losses [27, 28]. By identifying whether behavior in a gambling task is better explained by symmetric or asymmetric learning rules, this study addresses a core question about how value is updated in uncertain environments.

## 2 Results

### 2.1 Behavioral measures

47 participants (37 non-epileptic and 10 epileptic) performed a novel static risk-taking task (Starling task) in which they viewed one of two cards and decided whether it was higher (by pressing up-arrow on keyboard) or lower (by pressing down-arrow on keyboard) than an unseen opponent’s card, receiving trial-by-trial feedback. After the opponent’s card was revealed, participants learned whether their choice was correct. A correct response earned +$0.50, while an incorrect response resulted in a –$0.50 loss. The task had four blocks. The first block of cards was uniformly distributed from 1 to 9, and the second and third blocks were skewed low or high, with more or fewer presentations of lower and higher numbers, respectively (the blue box on the bottom in Fig. 1a illustrates the frequency of card numbers in each deck). These three blocks of trials in which the card distributions were fixed throughout the block, were followed by a mix block, in which all three distributions were interspersed, and the player was cued to which distribution deck the current card came from by the color of the card, which they previously learned in the fix blocks (see Section 4 Methods for full task details).

**Fig. 1.**
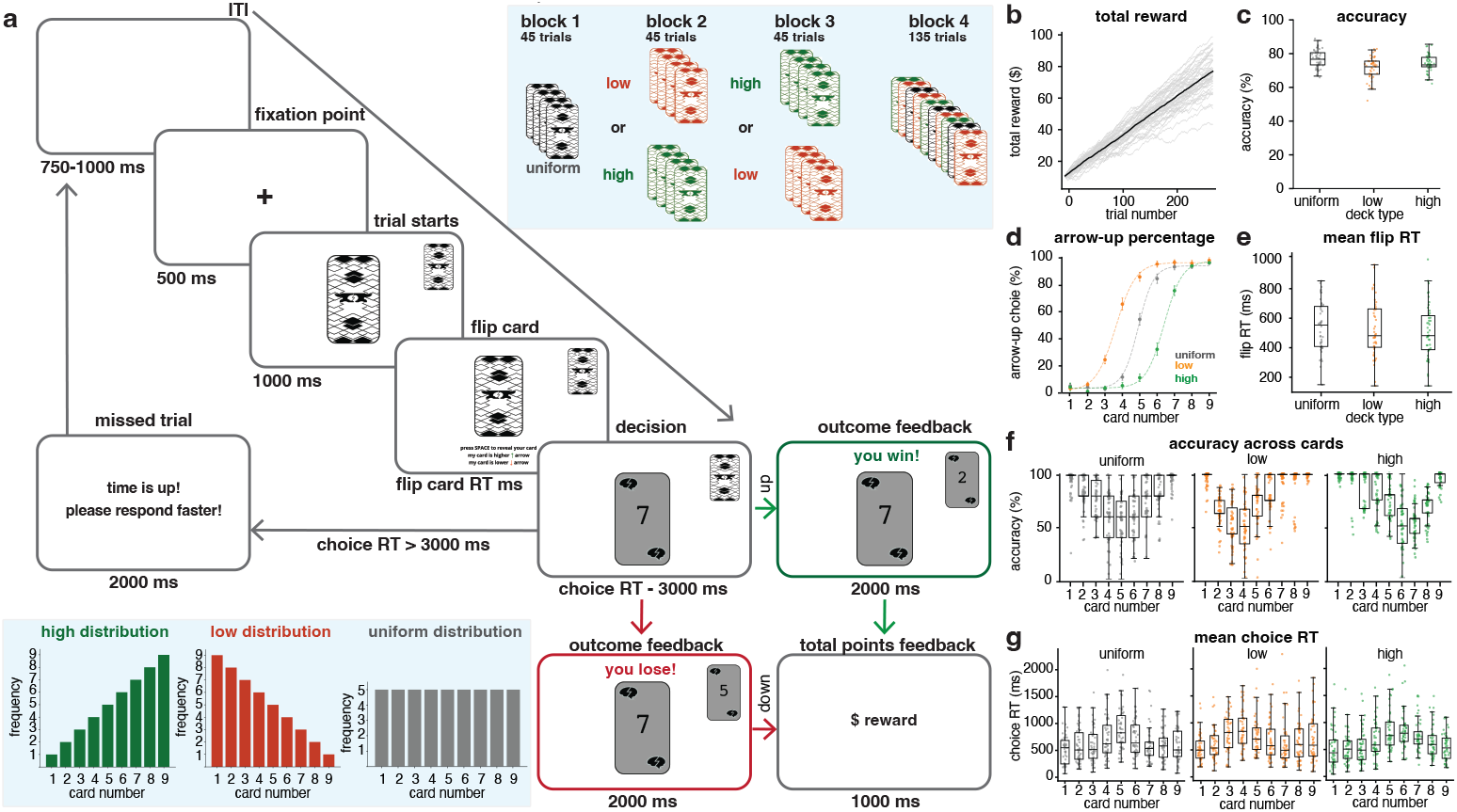
The Starling task and behavioral measures. **(a)** Task schematic. In each trial, participants flipped their card, decided if it was higher or lower than the opponent’s card, and received feedback. First 3 blocks are fix block and the last block is mix block (see blue box on top). **(b)** Participants’ total reward trajectories across trials in gray. The black line indicates the mean trajectory across participants. **(c)** Participants’ accuracy in blocks 1–3 for the uniform, low, and high decks. Each marker shows the mean accuracy across all trials for a single participant. **(d)** Mean percentage of arrow-up choices across participants in blocks 1–3, with error bars indicating SEM. Dashed curves represent the Sigmoid fits to the data points. **(e)** Flip RT distributions for each card deck and number in blocks 1–3. Each marker represents the average flip RT of a single participant. **(f)** Accuracy distributions for each deck and card number in blocks 1–3. Each marker represents the mean accuracy across all trials for a single participant. **(g)** Mean choice RT across participants and deck of cards in blocks 1–3. Each marker indicates the mean choice RT of a single participant.

Across trials, participants showed monotonic increases in total reward (Fig. 1b). Gray trajectories depict individual participants, and the black curve represents the across-participant mean. Mean accuracy by deck and block (Fig. 1c) was (M ± SEM (standard error of mean)%) across uniform (76.71 ± 0.7%), low (71.6 ± 0.9%), and high (74.3 ± 0.8%) card distributions. Choice behavior reflected the task structure: the percentage of arrow-up responses rose with card value and followed a Sigmoid curve (Fig. 1d), captured by three-parameter Sigmoid fits used in subsequent analyses. The arrow-down response curves exhibited a complementary pattern and are therefore not shown.

Combining across both non-epileptic and epileptic participants’ behavioral measures, mean flip RTs, which is defined as the interval between the appearance of the back of two cards and the participant flipping their own card, did not systematically vary by deck (Fig. 1e). Participants’ accuracies, defined as whether each decision was correct, varied systematically with card value, mirroring the task’s underlying uncertainty structure (Fig. 1f). Mean choice RTs, which were defined as the time elapsed between first card revealed and the participant’s decision (pressing arrow-up or arrow-down on the keyboard), were longest near the latent decision boundary (mid-range cards) and shorter for extreme card values (Fig. 1g).

### 2.2 Block structure modulates choice and timing

Behavioral measures were compared between blocks to study the fix (first 3 blocks - see Fig. 1a) and mix (last block - see Fig. 1a) blocks effects on participant choice and timing. Accuracy differed by block only in the low deck, with significantly lower performance in mix, 70.1 ± 1.2%, than in fix, 73.1 ± 1.0% (*t*(46) = 2.65, *p* = 0.03; Fig. 2a). Arrow-up choice probabilities and their Sigmoid fits for the two blocks are shown in Fig. 2b. Participant-level Sigmoid fits parameters (Fig. 2c) indicated that the midpoint (∼ 50% arrow-up) was displaced outward for the low/high decks in fix (low: 3.89 ± 0.14; high: 6.39 ± 0.09) but shifted toward the center (card 5) in mix (low: 4.37 ± 0.23; high: 6.24 ± 0.15), consistent with reduced reliance on extreme deck priors when distributions varied trial-by-trial. Slopes differed across decks between fix (low: 3.09 ± 0.25; high: 3.34 ± 0.24) and mix (low: 2.86 ± 0.28; high: 2.19 ± 0.25), whereas the uniform deck remained comparatively stable, suggesting altered choice sensitivity under contextual uncertainty (low: *Z* = −1.94, *p* = 0.04; high: *Z* = −2.98, *p* = 0.002). The outward midpoints in fix indicate reliance on deck base rates (priors), whereas the collapse toward the center in mix suggests reduced weighting of those priors when context varies on a trial-by-trial basis. The shift of midpoints toward the objective center (card 5) in mix shows reduced weighting of learned base rates in favor of trial-specific evidence (the revealed card value). This is consistent with base-rate neglect, which is the tendency to underweight prior probabilities when individuating information is available.

**Fig. 2.**
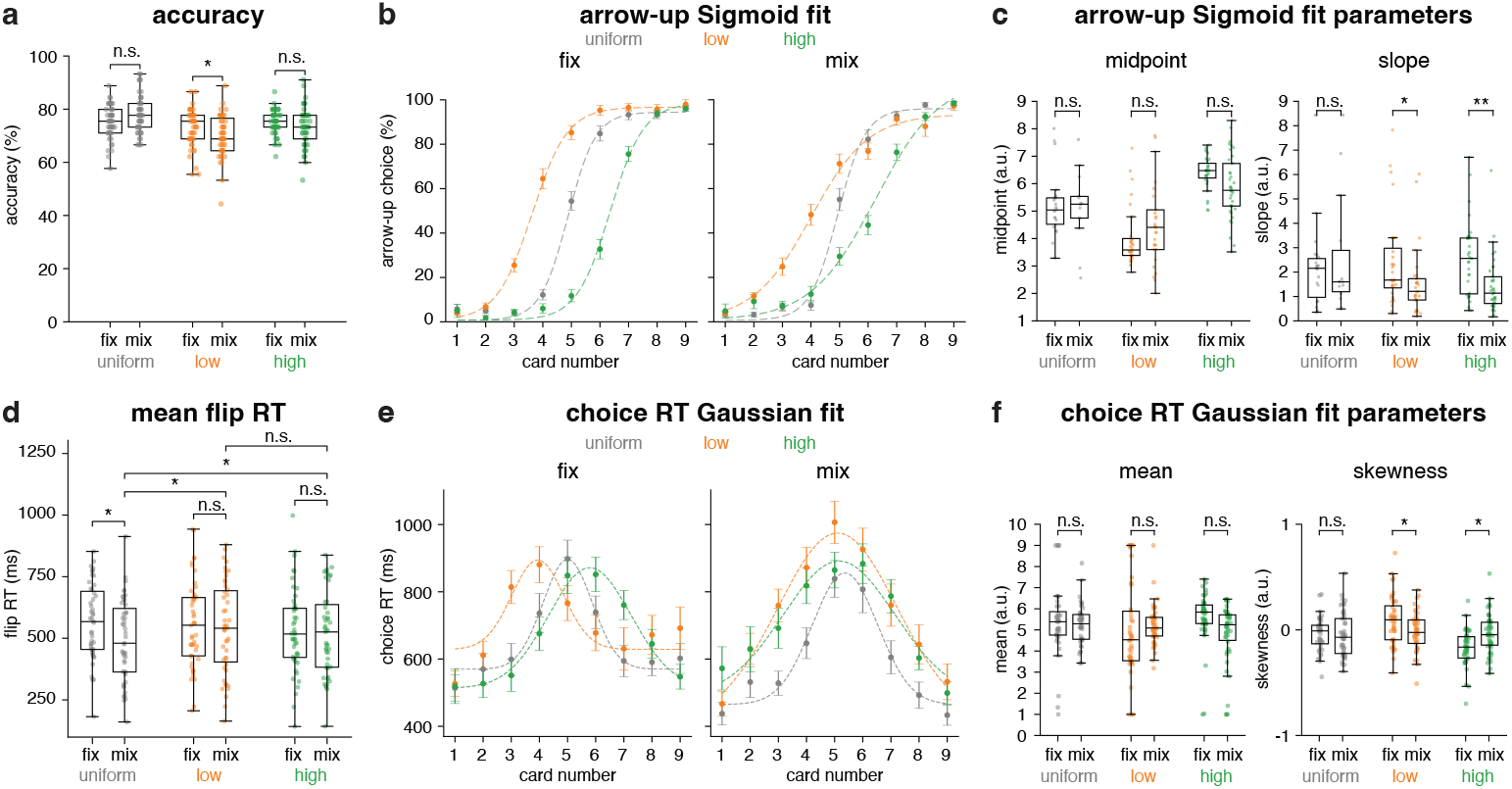
Behavioral measures in fix vs. mix blocks. **(a)** Accuracy (%) by block type and card decks. **(b)** Arrow-up choice (%) across card numbers for fix and mix blocks. Markers are mean percentage of arrow-up choices across participants, with error bars indicating SEM. Dashed curves represent Sigmoid fits to the data points. **(c)** Sigmoid fit parameters from (b) for each participant. **(d)** Flip RT distributions by card deck and number. **(e)** Mean choice RT distributions for each card and deck in fix and mix blocks, with error bars for variability. Dashed curves are Gaussian fits illustrating overall RT shapes. **(f)** Gaussian fit parameters from (e). Mean reflects central tendency and skewness shows RT asymmetry. In all panels, asterisks show fix vs. mix differences within decks (\**p* < 0.05, * * *p* < 0.01, n.s.= not significant).

Flip RTs were faster for the uniform deck in mix (492.51 ± 24.10 ms) than in fix (566.73 ± 22.15 ms; *t*(46) = 4.513, *p* < 0.001), with no reliable change for low or high decks (Fig. 2d). Furthermore, in the mix block flip RTs were faster for uniform deck compared to low (*t*(46) = −4.17, *p* < 0.001) and high (*t*(46) = −2.33, *p* = 0.025) decks in the same block (low: 541.79 ± 27.05 ms; high: 519.94 ± 24.77 ms). These results indicate that distributional uncertainty simplified decisions for uniformly distributed outcomes, whereas no such simplification was observed for skewed outcomes in the mix block. Mean choice RT functions across card values (Fig. 2e) were well summarized by Gaussian fits whose parameters revealed characteristic asymmetries (Fig. 2f). Choice RT distributions are shown as scatter–box plots in Fig. S1 (Supplementary Information). In fix blocks, low-deck RTs were right-skewed (mean: 4.92 ± 0.34; skewness: 0.09 ± 0.03) and high-deck RTs left-skewed (mean: 5.76 ± 0.17; skewness: −0.17 ± 0.02). In mix blocks, however, both low (mean: 5.18 ± 0.14; skewness: −0.01 ± 0.02) and high (mean: 4.82 ± 0.21; skewness: −0.04 ± 0.02) RT profiles converged toward the uniform pattern, indicating that trial-wise uncertainty collapsed deck asymmetries (low: *Z* = −2.22, *p* = 0.025; high: *Z* = 2.10, *p* = 0.034). The convergence of low/high RT asymmetries toward a more uniform RT profile in mix is consistent with the idea that participants rely less on deck-specific priors (base rates) and more on stimulus-driven difficulty that is similar across decks once context becomes volatile; i.e., uncertainty reduces the behavioral footprint of base-rate weighting in both choice functions and timing. This interpretation also aligns with accounts in which base-rate sensitivity depends on how participants represent the task and which strategy they retrieve under a given context [29, 30].

Together, these results show that the contextual uncertainty introduced in the mix block pulls the latent choice boundary toward the center, reduces slope differences across decks, and reshapes RT distributions toward a more uniform, ambiguity-driven profile. Relationships between accuracy and RTs in fix and mix blocks are illustrated in Fig. S2 (Supplementary Information). Fig. S2b shows that the fix block, which is less complicated, block accuracy decreases with longer choice RTs. This pattern does not reflect a classical speed–accuracy trade-off (SAT); instead, longer RTs likely show increased decision uncertainty that leads to both slower and less accurate responses. This interpretation is consistent with a study showing that speed and accuracy need not reflect a single built-in trade-off, but can dissociate depending on underlying strategic or task-related processes [31].

### 2.3 Preserved accuracy and slower choice RT in epilepsy

Because same candidate models were applied to both epileptic (N = 10) and non-epileptic (N = 37) participants, behavioral measures were compared between groups to determine whether any group differences could affect model fitting. Accuracy, flip RTs, and total reward did not show group differences (Fig. 3a,b,d). However, choice RTs were longer in the epileptic group (permutation; *p* < 0.05; Fig. 3c). Trial-wise trajectories further suggested group differences in block-related timing patterns (dashed gray lines indicate block onsets; Fig. 3e–f). Given the limited patient sample and potential non-task factors affecting RTs, the RT differences should be cautiously interpreted [32, 33].

**Fig. 3.**
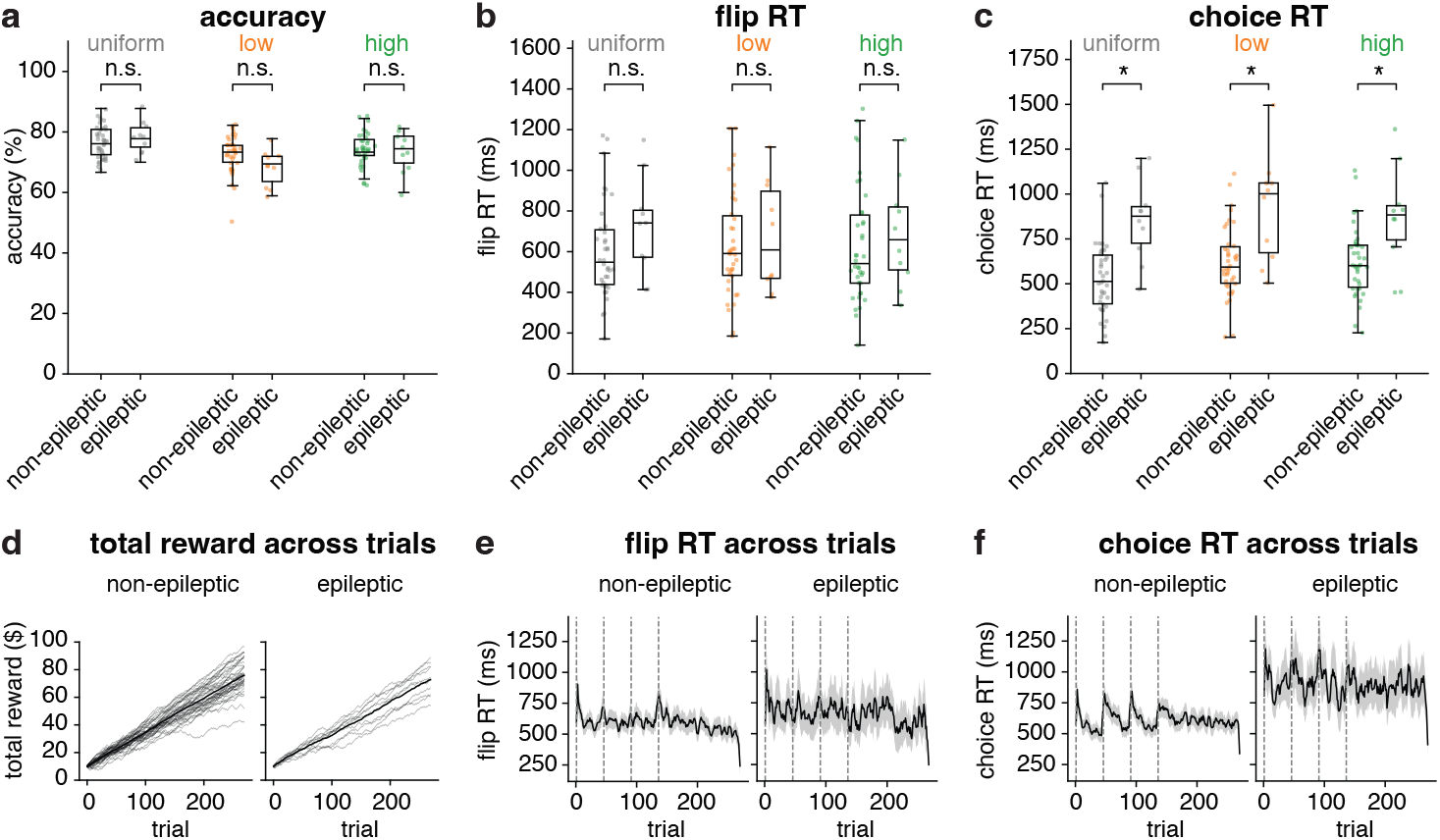
Comparison of behavioral performance between non-epileptic and epileptic participants. **(a)** Accuracy (%) did not differ significantly between groups. **(b)** Flip RTs showed no group differences. **(c)** Choice RTs were significantly longer in the epileptic group (asterisks indicate p < 0.05). **(d)** Trial-wise trajectories of total reward were not significant between the two groups (n.s., *p* > 0.05). **(e - f)** Trial-wise participants mean flip RT and mean choice RT, plotted separately for non-epileptic and epileptic participants. Solid curves denote mean RTs, and gray shaded bands indicate the SEM.

### 2.4 RS models perform Starling better than other candidate RL models

Before examining how the models mimic human behavior, their overall fit to participants’ behavioral data was first compared. Because the Starling task requires participants to learn action values from trial-by-trial feedback, behavior was modeled using RL frameworks that update expected values via RPEs and generate choices from those learned values on each trial. Five candidate RL models: win stay lose shift (WSLS), Rercorla Wagner (RW) with epsilon-Greedy, RW with Softmax, Dual–Q, and risk sensitive (RS) were evaluated on multiple metrics (see Section 4 Methods for detailed model specifications). Across accuracy, precision, recall, and specificity, which are metrics that quantify overall performance (see Section 4 Methods for more detail), RS achieved the highest mean scores (Fig. 4a). Significance testing of these metrics (Fig. 4b) showed RS significantly outperformed alternative models on accuracy, precision, recall, and specificity (paired-sample tests, *p* < 0.05).

**Fig. 4.**
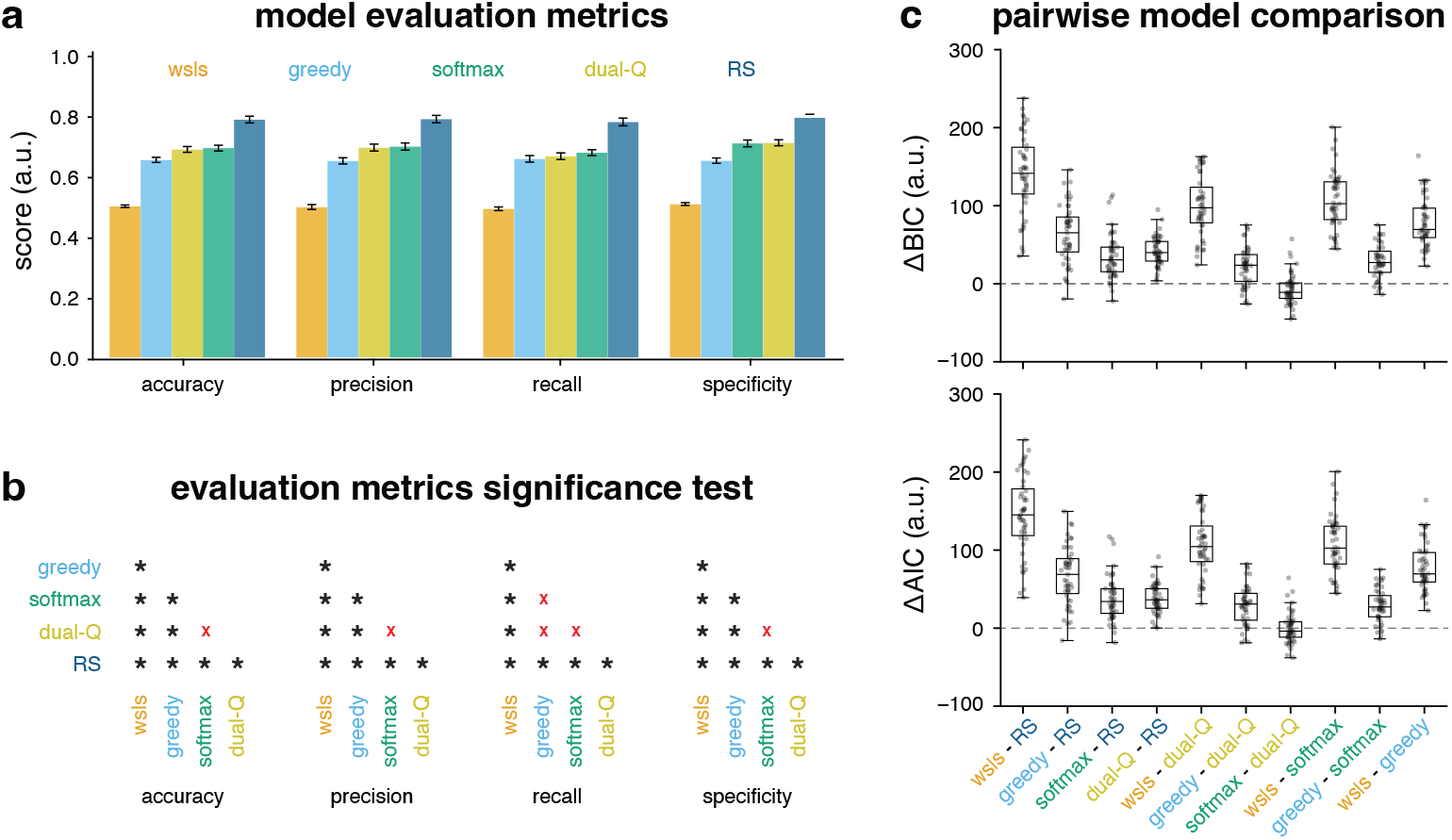
Evaluation and comparison of different learning models. **(a)** Performance metrics across candidate models. RS consistently outperformed ϵ-Greedy, Softmax, Dual-Q, and WSLS. **(b)** Pairwise significance testing of evaluation metrics. Asterisks indicate significant differences (p < 0.05), while red crosses denote non-significant results. **(c)** Pairwise model comparisons using ΔBIC ΔAIC.

Pairwise model selection via Bayesian Information Criteria (BIC) [34] and Akaike Information Criteria(AIC) [35] further favored RS model (Fig. 4c). Distributions of ΔBIC and ΔAIC were consistently shifted above zero for comparisons against RS, indicating a reliable preference for RS over ϵ-Greedy, Softmax, Dual–Q, and WSLS across participants. Overall, RS is the better-fitting model, whereas WSLS performs worst; therefore, WSLS was excluded from further analyses. Parameter recovery analyses supported interpretability of fitted parameters (Fig. S3).

### 2.5 RS aligns closely with human behavior

To test whether the models captured human behavior at the level of the task’s deck structure, model predictions were compared to human performance within each deck combining fix and mix blocks. Fig. 5a illustrates that RS did not differ significantly from participants in any deck, indicating the closest match to participants’ accuracy (paired t-test, asterisks: *p* < 0.05, n.s. not significant). Dual-Q showed significant deviations from participant behavior in most decks. Separated accuracies for fix and mix blocks are shown in Fig. S4 (Supplementary Information).

**Fig. 5.**
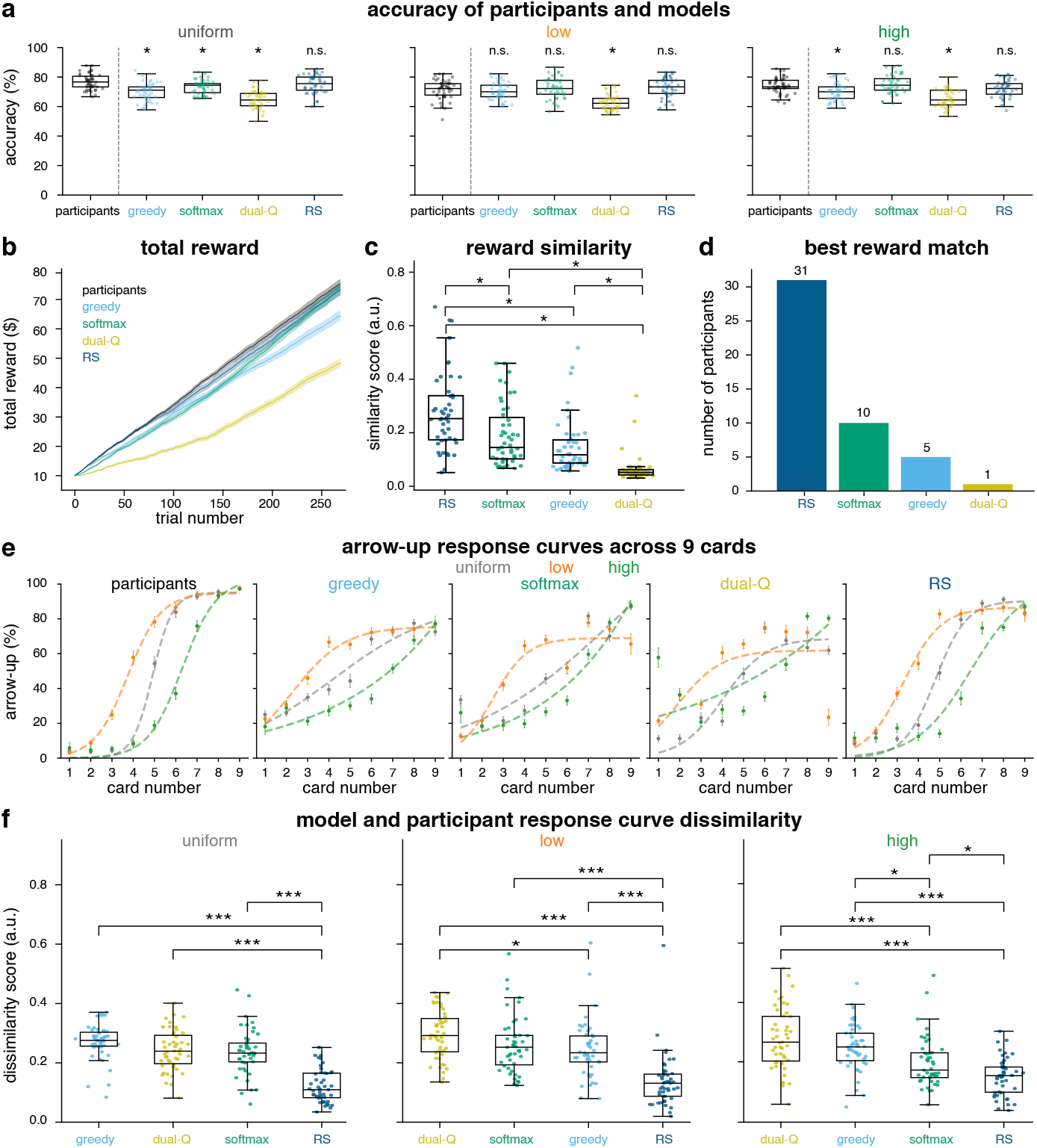
Model vs. participant behavior comparison. **(a)** Accuracy (%) across uniform, low, and high decks. Participants vs. models comparisons are displayed above each model panel (* *p* < 0.05, n.s: not significant). **(b)** Mean total reward trajectories throughout trials for each model. Solid curves denote the participants total reward mean, and shaded bands indicate the SEM. **(c)** Total reward similarity scores between participants and models. **(d)** Number of participants whose total reward trajectory is better explained by each model. **(e)** Arrow-up choice percentages Sigmoid fits across card deck and number, comparing participants with model predictions.**(f)** Arrow-up choice percentages Sigmoid fits dissimilarity scores between participants and models for each card decks. Asterisks show significant differences (\**p* < 0.05, * * *p* < 0.01, * * \**p* < 0.001)

Trial-wise reward trajectories revealed that RS most closely tracked participants’ reward trajectories (Fig. 5b). This was quantified with reward similarity scores in Fig. 5c (see Section 4 Methods), compared across models. RS scores exceeded those of Softmax, ϵ-Greedy, and Dual-Q (paired ANOVA followed by Bonferroni-corrected post hoc tests; asterisks: *p* < 0.05). Consistently, the number of participants better explained by each model (Fig. 5d) favored RS (31 participants) over Softmax (10), ϵ-Greedy (5), and Dual-Q (1), confirming RS as the most likely model of participants’ reward trajectories.

RS also reproduced the Sigmoidal choice functions observed in participants across uniform, low, and high decks, with sharper transitions between card values, consistent with more confident decisions at the extremes (Fig. 5e). By comparison, *ϵ*-Greedy responses appeared more threshold-like, Softmax smoother and less steep, and Dual-Q deviated most from participants’ empirical curves.

Dissimilarity between participant and model Sigmoid fits (see Section 4 Methods; Eq. 26) was compared within each deck (Fig. 5f). RS yielded the lowest dissimilarity in all decks (ANOVA followed by Bonferroni-corrected post hoc tests; * *p* < 0.05, * * *p* < 0.01, * * * *p* < 0.001). The same pattern held upon separately analyzing both the fix and mix blocks (Fig. S5, Supplementary Information): RS again exhibited the lowest dissimilarity of all models in both blocks.

RS consistently outperformed *ϵ*-Greedy, Softmax, and Dual-Q at accuracy matching, reward-trajectory similarity, and response-curve dissimilarity. Since RS, *ϵ*-Greedy, and Softmax all significantly outperformed the Dual-Q in all of these measures, the Dual-Q model was excluded from subsequent analyses. Across all behavioral benchmarks, RS showed the highest correspondence to participant performance, identifying it as the best behavioral model for the Starling task for most subjects.

These differences reflect distinct computational assumptions. *ϵ*-Greedy and Softmax assume symmetric value updating from gains and losses, differing only in how they implement exploration. Dual-Q separately tracks reward and risk signals. In contrast, RS updates values asymmetrically using separate learning rates for positive and negative RPEs. The results therefore suggest that most participants relied on asymmetric outcome updating rather than symmetric learning or separate reward–risk tracking.

### 2.6 Model-derived latent variables explain participant choice and timing

To understand why the models differ in their behavioral predictions, we examined the effect of trial-by-trial Q-values on behavior for each model. These Q-values represent learned reward estimates of action value that are updated from feedback and used to guide subsequent choices. (see Equation 4 in Section 4 Methods for more detail on Q-value). For each model, the mean Q-value difference ΔQ = Q_arrow-up_ − Q_arrow-down_ was plotted by card and deck (Fig. 6a). RS displayed the largest dynamic range with pronounced nonlinearity, yielding sharper transitions between adjacent card values, consistent with more confident decisions at the extremes. Softmax produced smoother, more gradually changing curves, whereas *ϵ*-Greedy appeared more threshold-like. These patterns indicate that RS-derived Q-values more closely reflect the task structure, particularly the amplified separation between high and low cards in skewed decks.

**Fig. 6.**
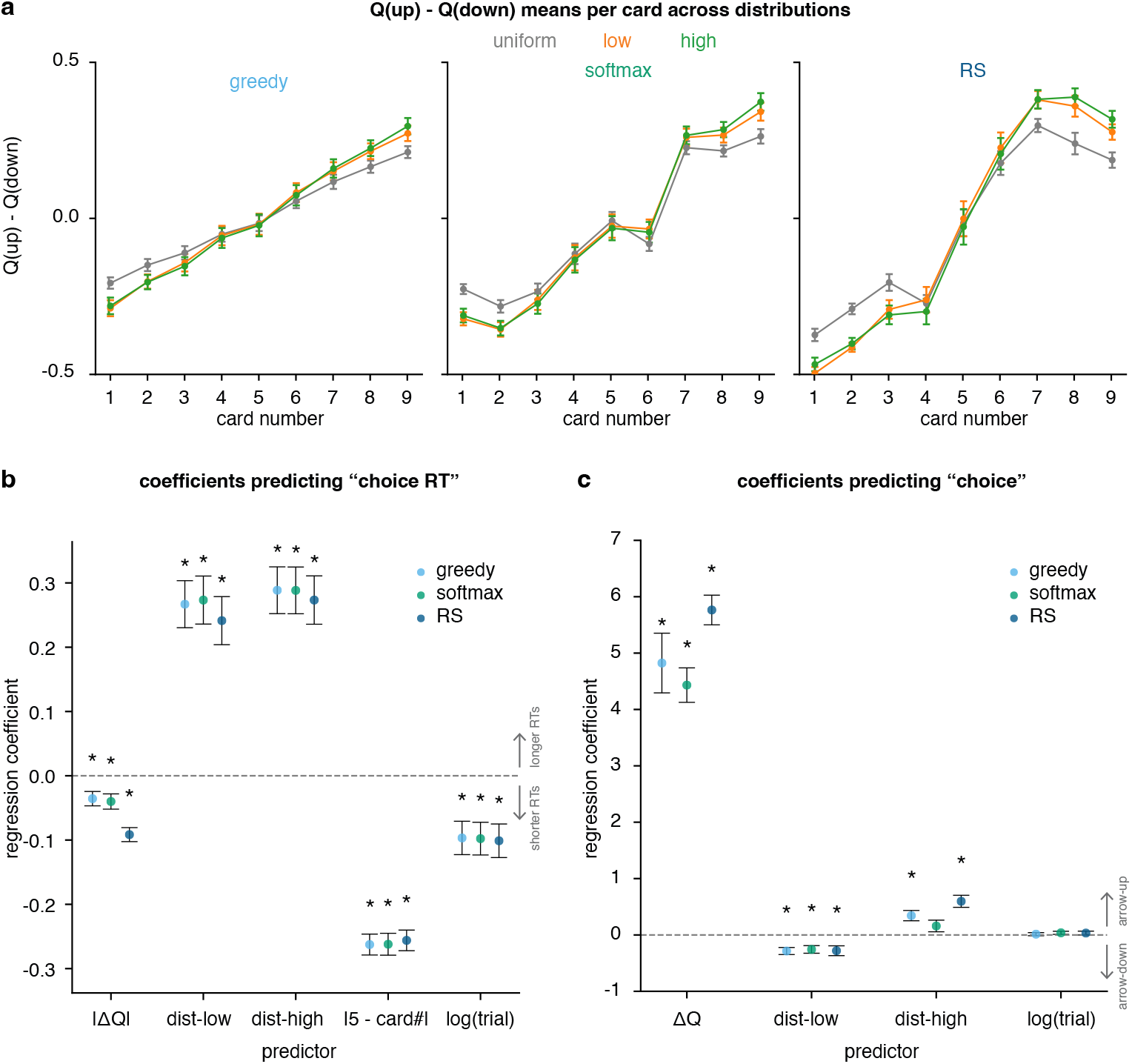
RL model variables’ association with choice probability. **(a)** Mean Q-value differences (Q_arrow-up_ − Q_arrow-down_) across card deck and number for candidate models. **(b)** Linear regression coefficients predicting choice RT from model-derived Q-values and task measures, for testing how different models explain subjects’ choice RTs. **(c)** Logistic regression coefficients predicting choices from model-derived Q-values and task measures, for testing how different models explain subjects’ choices. Asterisks denote significant effects (\**p* < 0.05).

To evaluate how model-derived value signals (Q-values) relate to choice RT, a linear regression was fit to predict trial-by-trial choice RT from both model-derived variables and task-related factors (Fig. 6b). Predictors included value separation (|ΔQ|), deck type, distance from the midpoint card (|5 − card#|), and trial number. This allowed us to assess the contribution of learned value signals in predicting the choice RT relative to structural features of the task. Regression coefficients quantify how changes in the predictors are associated with changes in RT. Statistical significance indicates whether each coefficient significantly differs from zero (Wald z-test with clustered standard errors, *p* < 0.05; see Table S4, Supplementary Information). Positive coefficients indicate slower responses (longer RTs), whereas negative coefficients indicate faster responses (shorter RTs). Near zero coefficients contribute less in the determining the choice RT.

Across models, |ΔQ| carried a significant negative coefficient, indicating that larger Q-value separations were associated with shorter RTs or faster decisions. Relative to the uniform baseline, low and high decks were associated with slower responses (positive coefficients). Distance from the midpoint card (|5 − card#|) carried a negative coefficient, indicating that responses were faster for more extreme card values (farther from 5). This pattern is consistent with lower decision uncertainty at the extremes, where the correct choice is more strongly constrained by the revealed card value. Trial number carried a negative coefficient, reflecting progressive speeding across task performance. Notably, although all models captured the difficulty effect, the RS model’s |ΔQ| showed the strongest (most negative) regression coefficient with choice RT. Regression analysis summary is provided in Table S4 (Supplementary Information). Together, these results show that trial-by-trial choice RT is shaped by model-derived value separation and task structure, with RS providing the most behaviorally predictive Q-value signal.

Participants’ binary choices (arrow-up vs. arrow-down) were regressed onto ΔQ to test whether model-derived values and task-related factors captured the computations guiding action selection on each trial. Predictors included ΔQ, deck type, and trial number. This allowed us to assess the contribution of learned value signals in predicting the choice relative to structural features of the task. Regression coefficients quantify how changes in the predictors are associated with changes in choice. In Fig 6c, logistic regressions on arrow-up/arrow-down choice identified ΔQ as the dominant positive predictor (Wald z-test with clustered standard errors, *p* < 0.05 -see table S5, Supplementary Information). Positive coefficients are associated with the arrow-up choice, while negative coefficients are associated with the arrow-down choice. Near zero coefficients contribute less to determining the choice.

Across all models, larger ΔQ indicated higher odds of choosing arrow-up (since ΔQ = Q_arrow-up_ − Q_arrow-down_). Deck indicators showed the expected directional effects relative to the uniform deck baseline: low deck carried a negative coefficient (participants were more likely to choose arrow-down since there are more low cards in that deck), and high deck carried a positive coefficient (participants were more likely to choose arrow-up since there are more high cards in that deck). The trial term did not significantly influence choice direction. The summary of these regression analyses is provided in Table S5 (Supplementary Information). Across value-difference profiles, RT regressions, and choice regressions, RS provided the most discriminative ΔQ signals, capturing both the nonlinearity induced by deck structure and the empirical link between value separation and response speed, thereby offering a better mechanistic account of both what participants chose and how quickly they decided.

In sum, trial-wise ΔQ robustly explained choice direction while RS yielded the clearest and most behaviorally aligned differences in value expectation across decks. Because participant choices remained probabilistic, even at large value separations, there exists an inherent noise ceiling on trial-wise predictability. While more flexible models could capture additional idiosyncratic variability, such gains would likely reflect overfitting rather than capturing task-relevant computations. The convergence of RS-derived ΔQ with both choice direction and response speed suggests that RS approaches this ceiling while retaining mechanistic interpretability.

## 3 Discussion

Across models, the RS model provided the best account of behavior, outperforming all alternative RL models on model-evaluation metrics and most closely reproducing key behavioral signatures of the Starling task. This result supports an asymmetric learning mechanism in which gains and losses update value at different rates, yielding an interpretable explanation for trial-by-trial choice patterns in environments with uncertain outcomes. Conceptually, asymmetric learning can be viewed as a simple approximation of distributional RL, in which different portions of the outcome distribution are updated at different rates. This may explain why RS performs well under uncertainty, as it captures sensitivity to distributional asymmetries without requiring an explicit representation of the full return distribution. This directly addresses our motivating question of whether symmetric updating is sufficient, or whether asymmetric learning from gains versus losses is necessary to explain trial-level choice under risk, and it provides interpretable latent variables. Notably, a small epileptic subgroup (10 epileptic and 37 non-epileptic) exhibited a comparable choice policy but slower RTs, suggesting that the computational mechanism of risk evaluation is preserved even when motor or processing speed is reduced [36]. Together, these findings position RS as a concise and transparent framework for computational decision-making under risk.

This study showed that choices in the Starling task, a novel static risk-estimation task, are best explained by an asymmetric RS learning rule. Across accuracy matching and behavioral dissimilarity measures, RS consistently outperformed RW with *ϵ*-Greedy, RW with Softmax, Dual-Q, and WSLS. Importantly, the RS model produced trial-wise value differences (ΔQ and |ΔQ|) that explained both *what* participants chose and *how fast* they decided. Together, these results suggest that dynamic choice behavior under risk is better captured by learning rules that weight rewards and losses differently than by symmetric updating alone. Across evaluation metrics, RS consistently outperformed the alternative models in accuracy, precision, recall, and specificity. This pattern aligns with prior work showing that risk-sensitive RL variants can outperform risk-neutral baselines when fitting human choice behavior [37].

Task manipulations further showed how contextual uncertainty changes the balance between prior information and trial-specific evidence. In fix blocks, midpoints shifted outward, consistent with stronger reliance on deck base rates (priors). In mix blocks, however, those midpoints moved back toward the center, suggesting that participants down-weighted deck priors when they had to consider trial-totrial variations in task context. This pattern matches classic base-rate neglect, in which prior probabilities are underweighted relative to case-specific evidence [38– 40]. Such reduced reliance on base rates is expected when prior information is less salient or perceived as less reliable [40]. Relatedly, recent work links individual differences in base-rate weighting to neural computations of prior expectations [41]. More broadly, reduced reliance on priors under changing or asymmetric evidence aligns with reports that people underweight rare evidence and show asymmetric decision biases in these settings [4].

Group comparisons also suggest that the computational strategy captured by RS generalizes across clinical status. Accuracy, flip RTs, and reward trajectories were comparable between epileptic and non-epileptic participants, while epilepsy was primarily associated with slower choice RTs. This dissociation is consistent with prior work showing that neurological or clinical factors can slow response execution without necessarily altering the underlying choice policy or learning computations [36]. In our data, the preservation of choice behavior alongside slowed decision times implies that the asymmetric value-updating mechanism identified here reflects a relatively stable decision computation, while timing may be more sensitive to non-specific factors such as processing speed, attention, or motor constraints. This separation has practical implications for future clinical and neural studies: it suggests that model-based comparisons of learning and choice mechanisms can remain informative even when overall response speed differs across groups, while emphasizing the need to treat RT-related effects as partially dissociable from choice policy.

Parameter recovery analyses indicated that model parameters were generally identifiable and interpretable across candidate models. For RS, α^+^, α^−^, and τ showed strong fitted–simulated agreement (reported Pearson correlation r, slope m, intercept b, and RMSE in Fig. S3a, Supplementary Information). Dual–Q parameters were also recoverable, although α_risk_ was estimated less precisely (Fig. S3b). Softmax and *ϵ*- Greedy exhibited good recovery profiles (Fig. S3c–d), whereas WSLS fits collapsed toward chance, indicating poor recoverability.

A notable feature of the RS model was that the loss learning rate α^−^ was estimated less precisely and was frequently near zero. When α^−^ ≈ 0, worse-than-expected outcomes generate only weak value updates, meaning losses are underweighted. As a result, options that occasionally yield large rewards retain relatively high Q-values, providing a mechanistic account of asymmetric sensitivity to gains versus losses. This pattern is consistent with evidence that apparent learning-rate asymmetries can partly reflect perseverative (choice-stickiness) tendencies [18]. Although the present α^−^ effect concerns choice learning, prior work shows that RPEs also shape memory, amplified by positive affect, raising the hypothesis that individuals with weaker loss learning may likewise show reduced memory for negative outcomes [12]. Parameter recovery used the same fitting pipeline and parameter bounds/grid resolution as the participant fits, ensuring comparability between simulated and empirical estimates.

There were several limitations in this study. First, although model selection converged strongly on RS, parameter precision (especially for α^−^) leaves room for hierarchical or fully Bayesian fitting to improve performance. Second, participant fatigue constrained the total number of trials, resulting in few observations of higher cards in the low deck and lower cards in the high deck (see Fig. 1a). Third, the overall sample size was modest (N = 37), with only 10 epileptic participants, which limits statistical power and constrains the generalizability of the findings. Replication in larger and more diverse samples will be important to confirm the robustness of these effects. Recent work shows meaningful between-participant heterogeneity in strategy use, including subgroups using dynamic arbitration, single-strategy policies, and different learning approaches; this cautions that a single better-fitting model at the group level can mask distinct individual solutions [2]. Consistent with this, developmental studies report substantial strategy heterogeneity (including non-learning/gambler’s-fallacy subsets) and more stochastic choice in adolescents [8].

Future work should broaden the modeling space and then link those models to neural data. On the modeling side, RS can be compared against richer sequence models (e.g., RNN [42, 43] or LSTM that carry trial history and latent context), additional RL families (model-based, Bayesian, hierarchical), and alternative policies (decaying *ϵ*-Greedy [6, 44], Thompson sampling [45], Upper Confidence Bound [46]). When the likelihoods of the observed choices given the model parameters, are intractable for these richer models, neural Bayes/simulation-based estimators (e.g., LaseNet) can recover trial-wise latent sequences from behavior, enabling robust model-based neural analyses [43]. On the neural side, trial-by-trial latent variables from each model (e.g., ΔQ, |ΔQ|, RPE^−^, RPE^+^, uncertainty/entropy) can be entered into time-resolved encoding models of iEEG spectral power at cue, choice, and feedback. Prior fMRI work reports ventral striatal BOLD correlates of RS RPE and Q-values, supporting neural plausibility of RS RL signals [14, 37].

## 4 Methods

### 4.1 Participants

A total of 47 individuals participated in the study, including 37 non-epileptic volunteers and 10 patients with drug-resistant epilepsy. The participants demographics summary is provided in Table S6 (Supplementary Information). Non-epileptic participants completed the task online through the laboratory website, while patients performed the same task via an offline browser in the Neuro Acute Care Unit at the University of Utah Hospital.

### 4.2 Task

The Starling task is a novel decision-making paradigm developed in our lab to investigate how people make value-based choices under uncertainty. Each trial consists of a sequence of timed events. Each trial began with an inter-trial interval (ITI) consisting of a blank screen presented for 750-1000 ms. A fixation cross was then displayed for 500 ms. Next, the backs of both participant’s and their opponent’s cards were shown for 1000 ms. An instruction message followed and remained on the screen until participants pressed the SPACE key to continue. After the SPACE press, a 500 ms delay preceded the card flip and choice window, during which the participant’s own card was revealed while the opponent’s card remained hidden; participants had up to 3000 ms to respond. Outcome feedback (win/lose or a missed-trial message) was presented for 2000 ms right after participant pressed arrow-up/arrow-down to respond. Finally, reward feedback was displayed for 1000 ms.

Card values ranged from 1 to 9. On each trial, both cards were drawn from the same deck type without replacement (i.e., from a finite set of cards) with the constraint that the two card values never matched. Decks were visually coded by color to indicate their underlying statistical distribution: the orange (low) deck was skewed toward low numbers (e.g., 1-3), the green (high) deck was skewed toward high numbers (e.g., 7-9), and the gray (uniform) deck assigned equal probability to all values (1-9).

Participants flipped and viewed their own card in order to estimate the likelihood that their card was higher or lower than the opponent’s based on both the revealed number and the statistical distribution of the deck. Participants responded by pressing the up arrow if they believed their card was higher, or the down arrow if they believed it was lower. If no response was made within 3000 ms, the trial was aborted and was repeated at the end of the block. Feedback was performance-based: each correct decision earned a hypothetical reward of $0.50, whereas each incorrect decision incurred a hypothetical penalty of $0.50. All participants began with a starting score of $10 points.

To experimentally manipulate uncertainty, the deck distributions were presented under two different types of task blocks: in the fix blocks, the deck type remained constant and was explicitly instructed at the start of each block. The block order was always uniform first, followed by low/high and high/low (counterbalanced across participants), to ensure that all participants first learned the baseline mapping under a stable, unbiased distribution before encountering the skewed distributions, while controlling for order effects between the low and high decks. In the mix block the deck distribution changed across each trial, and participants used the color cue to identify the deck distribution. Although the cue was fully observable, the mix block required participants to use the cue to adjust their behavior for the corresponding distribution on each trial.

Each participant completed 45 trials per condition (uniform, high, low), followed by 135 trials in the mix block in which deck distributions were randomized, totaling 270 trials. This design allowed for precise modeling of how participants incorporated both outcome distributions and uncertainty into their decisions. It also enabled us to test various RL models against human behavior under varying probabilistic contexts.

The task was implemented and run online using jsPsych [47]. This browserbased implementation made the paradigm portable, standardized, and reproducible, with automated capture of trial-by-trial choices, RTs, and outcomes under identical stimulus timing and response rules across participants. Importantly, the full task is available for reuse, enabling other researchers to replicate the experiment directly or adapt it for related studies of decision-making under uncertainty. See Declaration for more information about task and related codes.

### 4.3 Computational models

Five common RL-based computational models were trained to capture participants’ choice behavior. The first model, Win–Stay Lose–Shift (WSLS), implements the simplest heuristic strategy. All remaining models were based on a Rescorla–Wagner (RW) RL framework. Among these, one model employed a Softmax policy for probabilistic action selection, while another used a *ϵ*-Greedy policy. Two additional models, referred to as the dual–Q and RS, extended the RW framework to include separate learning rates for positive and negative outcomes allowing asymmetric learning from rewards and losses.

#### 4.3.1 Win stay lose shift (WSLS) model

The WSLS heuristic models choice behavior as a simple function of the previous outcome. The model is parameterized by two probabilities: the probability of staying with the same action after a win, *p*(stay | win), and the probability of shifting after a loss, *p*(shift | loss). Formally, the probability of the current action *a*_*t*_ given the previous action *a*_*t*−1_ and reward *r*_*t*−1_ is

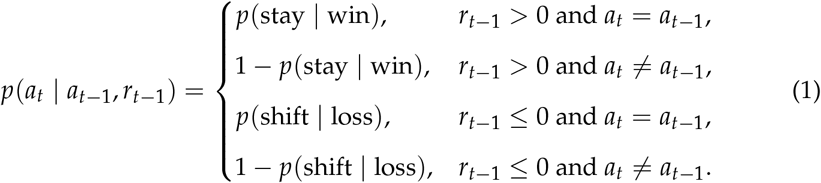

Thus, after a rewarded trial the model tends to repeat the same action with probability *p*(stay | win), while after an unrewarded trial it tends to switch the action with probability *p*(shift | loss). For the grid search process, the stay and shift probabilities were each sampled once from a uniform distribution *U* at the start of model fitting:

For *N* = 100 independent samples, *i* = 1, …, 100 :

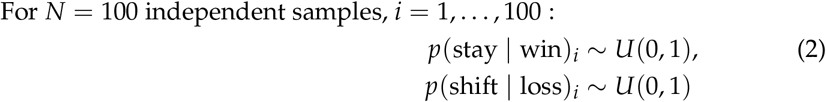

Accordingly, each of the two free parameters was sampled at 100 values, producing a total of 10,000 candidate parameter pairs evaluated during grid search.

#### 4.3.2 Rescorla-Wagner framework

In model fitting process, the RW agent was trained to predict trial-by-trial choice behavior for each participant. Q-values were initialized for every card number, distribution, and action as small random values close to zero.

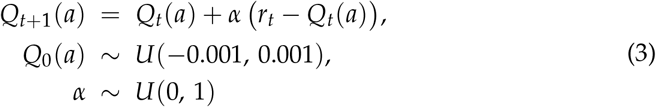

where *Q*_*t*_(*a*) is the expected value of choice *a* at trial *t, α* is the learning rate, *r*_*t*_ is the –$0.50 or +$0.50 reward at trial *t*, and *U* is a Uniform distribution. Each RW model maintains a three-dimensional Q-table across card numbers, deck distributions, and actions:

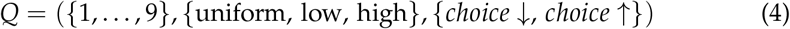

The Q-values were then translated into choice probabilities to determine action selection. Action selection is governed by different policies and the fitting involves finding the parameters that explain participant choices better. All parameters in each model were estimated by grid search over uniformly spaced values. Parameter values were sampled once at the beginning of each training process before starting the iterations. The best parameters for different models were estimated by computing the log-likelihood of observed choices under those parameters. The parameter set that maximized log-likelihood was selected as the best fit. Standard guidelines was followed for computational modeling, framing models to link observable variables to behavior and using simulation, parameter estimation, model comparison, and latent-variable inference to relate models to data [48]. In addition, multiple RL model families were compared instead of presupposing a single model, as the optimal strategy depends on environmental structure [49].

#### 4.3.3 Epsilon-Greedy model

In this model, while using RW method as Q-value update rule, at each trial, the agent chooses either to exploit the action with the highest estimated value or to explore alternative actions with a small probability. The choice probability for action *a* is defined as:

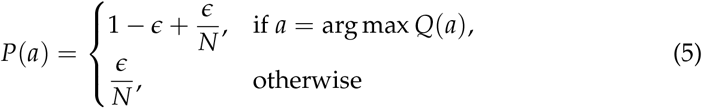

where *ϵ* is the exploration parameter and *N* is the number of available actions. With probability (1 − *ϵ*), the agent exploits by selecting the action with the highest Q-value, and with probability *ϵ*, it explores by choosing randomly among all other actions. To estimate these parameters, samples from the Uniform distribution *𝒰* were taken once at the start of model fitting:

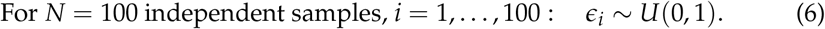

For each participant, the log-likelihood of the observed choice sequence was evaluated across all 10,000 sampled (*α, ϵ*) pairs (100 values per parameter), and the pair that maximized the log-likelihood was selected as the best-fitting parameter set.

#### 4.3.4 Softmax model

In this model, while using RW method as Q-value update rule at each trial, the probability of selecting action *a*_*i*_ is given by:

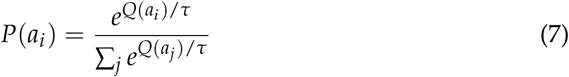

where *Q*(*a*_*i*_) is the estimated value of action *a*_*i*_, and *τ* is the temperature parameter controlling choice stochasticity. Lower values of *τ* make the model more deterministic and greedy, favoring actions with higher Q-values, whereas larger *τ* increases randomness, producing more exploratory behavior. To estimate the model parameters, *α* and *τ* were sampled from a Uniform distribution *𝒰* at the start of model fitting:

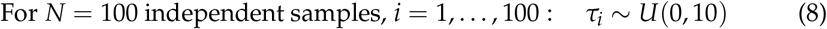

Accordingly, for each participant, the log-likelihood of the observed choice sequence was evaluated over 10,000 candidate (*α, τ*) combinations (100 sampled values for each parameter), and the combination yielding the maximum log-likelihood was selected.

#### 4.3.5 Risk-sensitive (RS) model

The RS model extends the RW framework by incorporating separate learning rates for positive and negative RPE, allowing asymmetric updating based on whether an outcome is rewarding or punishing. This asymmetry captures participants’ potential sensitivity for reward and loss.

At each trial, the action value *Q*(*a*_*t*_) is updated according to the RPE *δ*_*t*_ = *r*_*t*_ − *Q*_*t*_(*a*_*t*_):

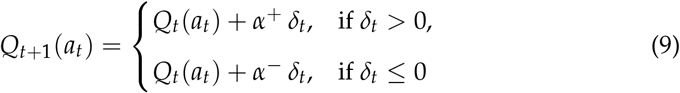

where *α*^+^ and *α*^−^ denote the learning rates for positive and negative RPE, respectively. A larger *α*^+^ indicates faster updating following rewarding outcomes, whereas a larger *α*− reflects stronger sensitivity to losses. Action selection follows a Softmax policy based on the current Q-values. This asymmetric update is a special case of RS where a nonlinear transform of the RPE induces gain/loss asymmetries; see Shen et al. for a general framework and convergence guarantees [37]. In experience-based risky choice, differential learning from gains versus losses, together with nonlinear utility, predicts risk preferences, supporting the use of asymmetric learning rates here [15].

Model parameters were sampled from a Uniform distribution *U* once at the start of model fitting:

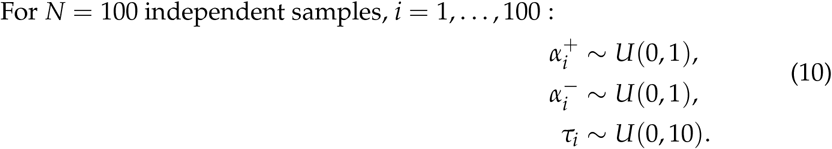

For each participant, the log-likelihood was computed over the full grid of 1,000,000 (*α*^+^, *α*^−^, *τ*) combinations (100 sampled values per parameter), and the maximizing triplet was retained as the best fit.

#### 4.3.6 Dual–Q model

The Dual–Q model extends the RW learning rule by maintaining two independent value functions, one tracking reward expectancy (Q^reward^) and another tracking risk expectancy (Q^risk^). This allows the agent to learn separately from rewarding and aversive outcomes.

At each trial, the model computes the risk associated with the shown card1 as the uncertainty of whether the second not shown card2 will be higher or lower:

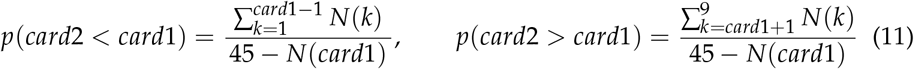

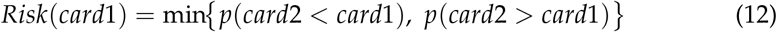

where *N(k)* is the number of remaining cards of value *k*. Cards with a similar probability of being higher or lower are considered riskier (less certain). All risks calculated using the equation above for uniform, low, and high decks are presented in Tables S1–S3.

The two value functions are then updated using separate learning rates according to the received reward *r*_*t*_ and estimated risk Risk(card1_t_):

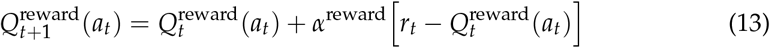

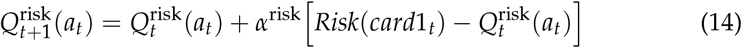

The combined value guiding choice behavior is then computed as:

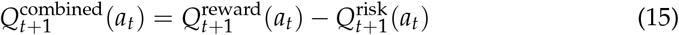

The combined value 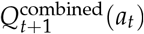 was then used in the Softmax policy to compute action-selection probabilities, encouraging choices that maximize expected reward while minimizing expected risk.

Model parameters were sampled from a Uniform distribution *𝒰* once at the start of model fitting:

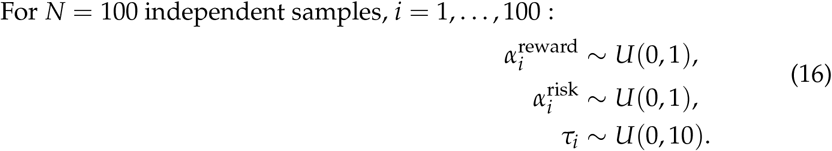

For each participant, the log-likelihood was computed over the full grid of 1,000,000 (*α*^reward^, *α*^risk^, *τ*) combinations, and the maximizing parameter set was retained as the best-fitting solution.

### 4.4 Behavioral analysis

Several behavioral measures were extracted from the task for use in both behavioral and modeling analyses. Accuracy was defined as the proportion of correct trials (e.g., choosing “arrow-down” when the participant’s card was 2 and the opponent’s card was 6, or choosing “arrow-up” when the opponent’s card was 1). Total reward was defined as the cumulative point total across trials. Choice preference was quantified as the proportion of “arrow-up” (participant’s card is higher) versus “arrow-down” (participant’s card is lower) responses. Flip RT was defined as the response time from the onset of the backs of the cards to the participant pressing the SPACE key to reveal their card; flip RTs exceeding ± 1.5× the standard deviation from each participant’s mean flip RT were excluded. Choice RT was defined as the response time from card reveal to the participant’s decision (pressing the up or down arrow key).

#### 4.4.1 Sigmoid fits to arrow-up percentage distributions

The parameters (*L, x_0_, k*) were estimated by fitting the Sigmoid to arrow-up percentages across all 9 card values, separately for the uniform, low, and high deck distributions. Three-parameter Sigmoid functions were fit to the proportion of arrow-up responses across card numbers (Figs. 1d, Fig. 2b, and Fig. 5e):

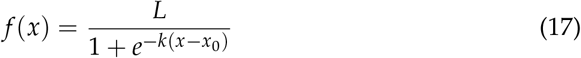

where *L* denotes the upper asymptote, *x*_0_ is the midpoint at which *f(x_0_)* = *L*/2, and k is the slope parameter that determines the steepness of the curve.

#### 4.4.2 Gaussian fits to choice RT distributions

To characterize the shape of choice RT distributions, Gaussian functions were fitted to mean choice RT values across card numbers, separately for each deck type (uniform, low, high) and block (fix, mix; Fig. 2e). The fitted function was:

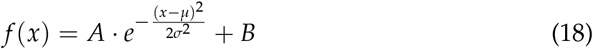

where *A* is the amplitude, *µ* is the mean, *σ* is the standard deviation, and *B* is a baseline offset. In addition, the asymmetry of the distributions was quantified by computing the skewness of the fitted curves:

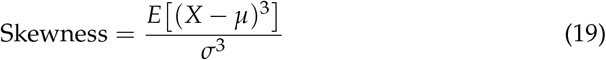

where *E*[·] denotes the expectation over the distribution.

### 4.5 Statistical analysis

Unless stated otherwise, all behavioral and model-performance statistics were conducted on participant-level summary measures. All tests were two-tailed and used *α* = 0.05.

#### Fix vs mix comparisons (Fig. 2)

Accuracy and mean flip RT were compared between fix and mix blocks using paired-sample *t*-tests within participants, performed separately for each deck distribution (uniform/low/high; Fig. 2a,d). To control the family-wise error rate across the three distribution-specific comparisons for each outcome, *p*-values were Bonferroni-adjusted (*m* = 3). For fitted curve parameters (Fig. 2c,f), fix vs mix differences were assessed using the Wilcoxon signed-rank test due to non-normality of parameter distributions.

#### Correlation analyses (Fig. S2)

Within each block (fix and mix), associations between participants’ mean flip time, choice time, and accuracy were evaluated using Pearson correlation. Statistical significance was assessed using permutation tests by randomly permuting participant labels 10,000 times to generate a null distribution of correlation coefficients. Two-tailed *p*-values were computed as the proportion of permuted correlations with absolute value greater than or equal to the observed correlation.

#### Group comparisons (Fig. 3)

Behavioral measures were compared between the non-epileptic and epileptic groups using permutation tests (10,000 iterations). Two-tailed *p*-values were computed as the proportion of permuted group differences whose absolute value was greater than or equal to the observed difference. For each behavioral outcome, *p*-values were Bonferroni-adjusted across the three distributions (*m* = 3).

#### Model comparison on classification metrics (Fig. 4b)

For each participant and model, binary class predictions (arrow-up = 1, arrow-down = 0) were generated by sampling from the model’s Softmax probabilities on each trial. Predicted choices were compared against participants’ observed choices to construct a confusion matrix with true positives (TP), true negatives (TN), false positives (FP), and false negatives (FN).

Classification metrics were defined as:

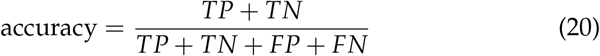

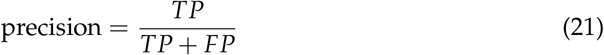

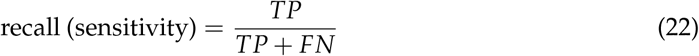

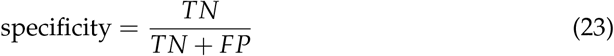

Accuracy reflects how often the model and participant made the same choice. Precision indicates how often the model’s arrow-up predictions were correct. Recall (sensitivity) measures how well the model captured participant arrow-up choices. Specificity measures how well the model captured participant arrow-down choices.

Models were compared using paired-sample tests across participants separately for each metric. Pairwise model comparisons were Bonferroni-adjusted within each metric for the number of model-pair tests performed (i.e., 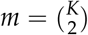 for K models).

#### Model vs participant accuracy (Fig. 5)

Participant accuracy was compared against each model’s predicted accuracy using paired-sample tests across participants (participant minus model), with multiplicity controlled as described above. Differences among the three deck distributions within a given block/condition (panels c and f) were tested using a one-way repeated-measures ANOVA with distribution (uniform/low/high) as a within-subject factor. When the omnibus ANOVA was significant, post-hoc paired comparisons between distributions were conducted with multiplicity control within the post-hoc family.

#### Regression analyses (Fig. 6)

Regression coefficients were tested using two-tailed Wald *z*-tests with participant-level cluster-robust standard errors (panels b and c). For each regression family (i.e., set of coefficients tested within a panel), *p*-values were corrected for multiple comparisons using Bonferroni adjustment. Full regression outputs and corrected *p*-values are provided in Tables S4-S5.

### 4.6 Models performance evaluation

#### 4.6.1 Models vs. human

In Fig. 5c for each participant and model, a fit score was computed to quantify how closely the model-predicted total reward trajectory matched the participant’s observed total reward. The score was defined as the absolute value of the Pearson correlation coefficient between the participant and model reward trajectories, divided by the root mean squared error (RMSE):

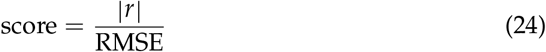

This metric jointly captures the similarity between the total reward trajectories of participants and models. The resulting scores were compared across models using a one-way ANOVA. In Fig. 5d, for each participant, the model yielding the highest score was identified, and the total number of participants best fit by each model was summarized in a histogram.

In Fig. 5f to assess model–participant dissimilarity in Sigmoid fits (Fig. 5e), the fitted parameters vectors were compared using the Euclidean distance:

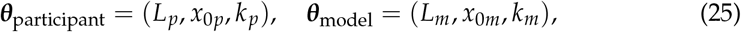

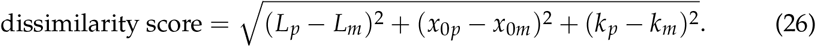

The dissimilarity scores in Fig. 5f were compared across models using a one-way ANOVA to test for overall differences.

#### 4.6.2 Regression

In Fig. 6, Logistic and linear regression models were used to examine which model parameters predicted participants’ choices and choices RTs, respectively.

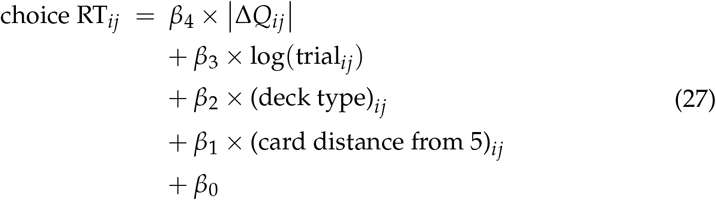

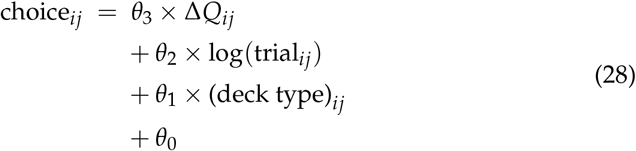

where choice_*ij*_ and choice RT_*ij*_ denote the binary choice outcome and RT for participant *i* and trial *j* ; Δ*Q*_*ij*_ is the difference in model-estimated Q-values between the two decks; |Δ*Q*_*ij*_| is its absolute value, indexing decision difficulty; trial_*ij*_ is trial *j* for participant *i* (log-transformed to capture learning or fatigue effects); (deck type)_*ij*_ indicates the deck distribution context (uniform, low, and high) dummy-coded with uniform as the reference level such that the coefficients for deck-low and deck-high represent differences in mean choice or RT relative to the uniform condition; (card distance from 5)_*ij*_ represents the absolute difference between the displayed card and card number 5; and *β*_*k*_, *k* = 1, 2, 3, 4 and *θ* _l_, l = 1, 2, 3 are regression coefficients estimating the influence of each predictor, with *β*_0_ and *θ*_0_ as intercepts.

In each linear and logistic regression model clustered standard errors was used at the participant level to account for within-subject correlations across trials. Because repeated trials from the same participant are not independent, all standard errors are computed using cluster-robust variance estimation. Here the cluster is the participant *i*. This correction inflates the standard errors to account for within-participant correlation, ensuring that *p*-values and confidence intervals remain valid even though each participant contributes multiple correlated trials [50–52]. To control for multiple hypothesis testing across predictors, *p*-values are adjusted using Bonferroni correction [53].

## Declarations

### Supplementary information

included

### Ethics approval and consent to participate

all participants provided informed consent in accordance with institutional review board guidelines. Both online participants and epilepsy patients provided informed consent before completing the task.

### Online task

https://www.neurosmiths.org/

### Code and data

OSF-Static Learning Risk Task (Starling) or GitHub

### Author contributions

E.S., R.C., A.B., and N.S. designed the experiment and N.S implemented the experiment using jsPsych. N.S., R.C., and A.P. conducted the experiments and collected data. N.S. performed the analyses. A.L. contributed to the model design. During the design and implementation of the online task, R.R., V.Z., and M.L. participated in regular pilot testing and provided feedback. S.R. and B.S. performed the surgical implantations and facilitated patient participation and task administration. T.D. provided technical and IT support. All authors discussed the results and reviewed the manuscript.

