## supplementary information for "Asymmetric Reinforcement Learning Explains Human Choice Patterns in Decision-making Under Risk"

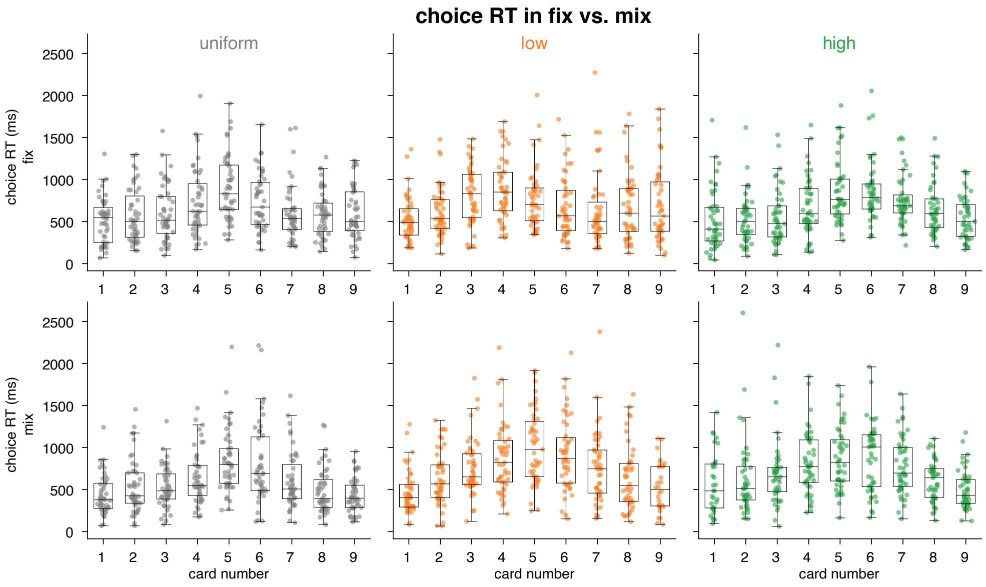


**Fig. S1** Mean choice RT across participants and deck of cards for fix blocks (first row) and mix blocks (second row). Each marker indicates the mean choice RT of a single participant in each category.

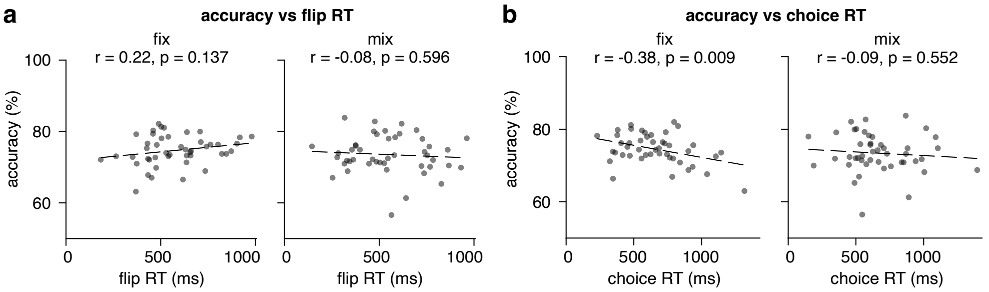


**Fig. S2** Relationship between accuracy and the two RTs in fix and mix blocks. (**a**) Accuracy vs. flip RT in fix and mix blocks. Accuracy shows no reliable association with flip RT in either block. (**b**) Accuracy vs. choice RT in fix and mix blocks. In the fix block (the block with less uncertainty) accuracy decreases with longer choice RTs that shows the speed accuracy trade-off. Correlation coefficients (*r*) and *p*-values are shown in each panel.

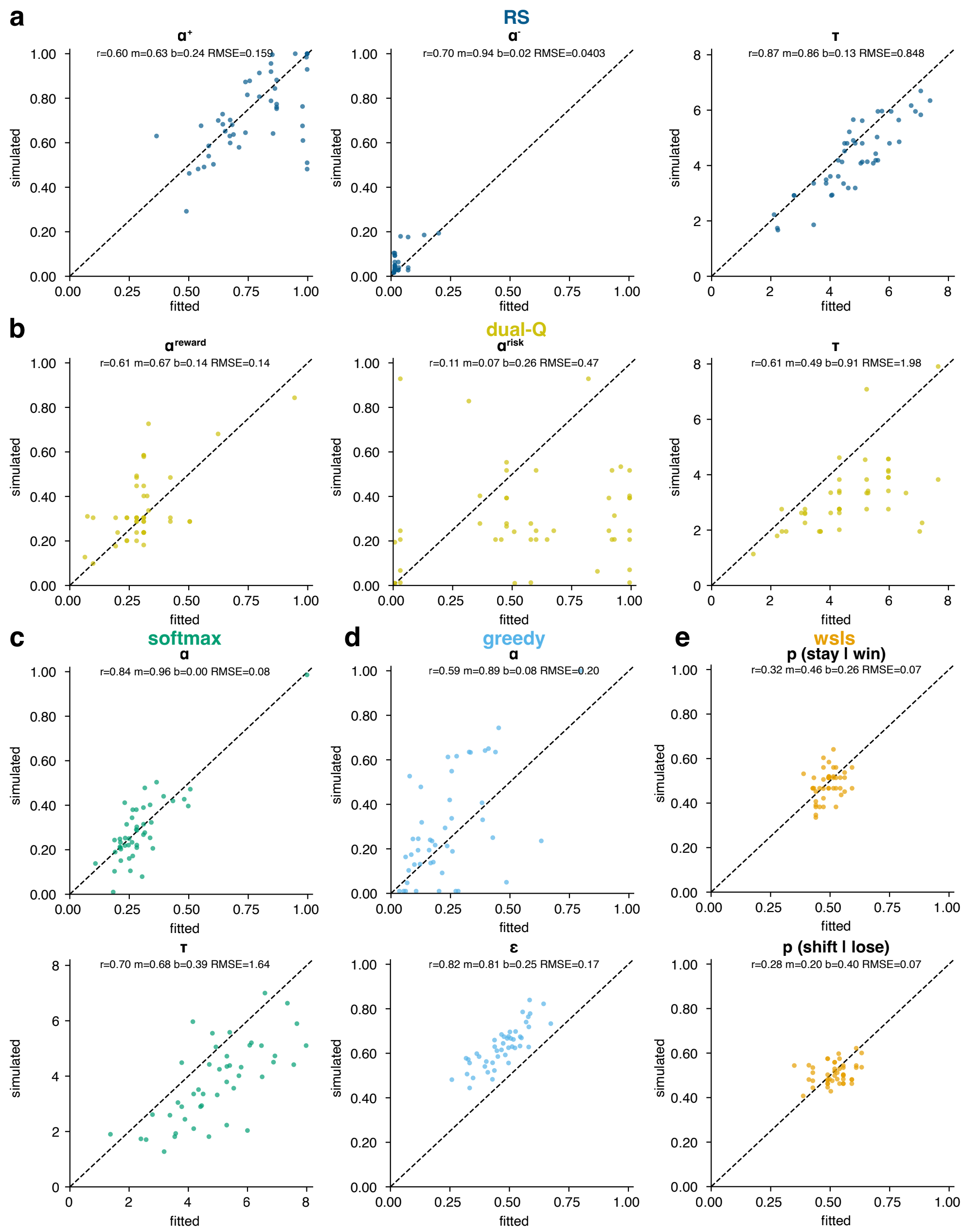


**Fig. S3** Parameter recovery for models. Scatterplots show fitted vs. simulated values with recovery metrics (Pearson correlation $r$, slope $m$, intercept $b$, and Root Mean Squared Error). (a) RS parameters: $\alpha^+$, $\alpha^-$, $\tau$. (b) Dual-Q parameters: $\alpha_{reward}$, $\alpha_{risk}$, $\tau$. (c) Softmax parameters: $\alpha$, $\tau$. (d) Greedy parameters: $\alpha$, $\epsilon$. (e) WSLS parameters: $p(stay|win)$, $p(shift|lose)$.

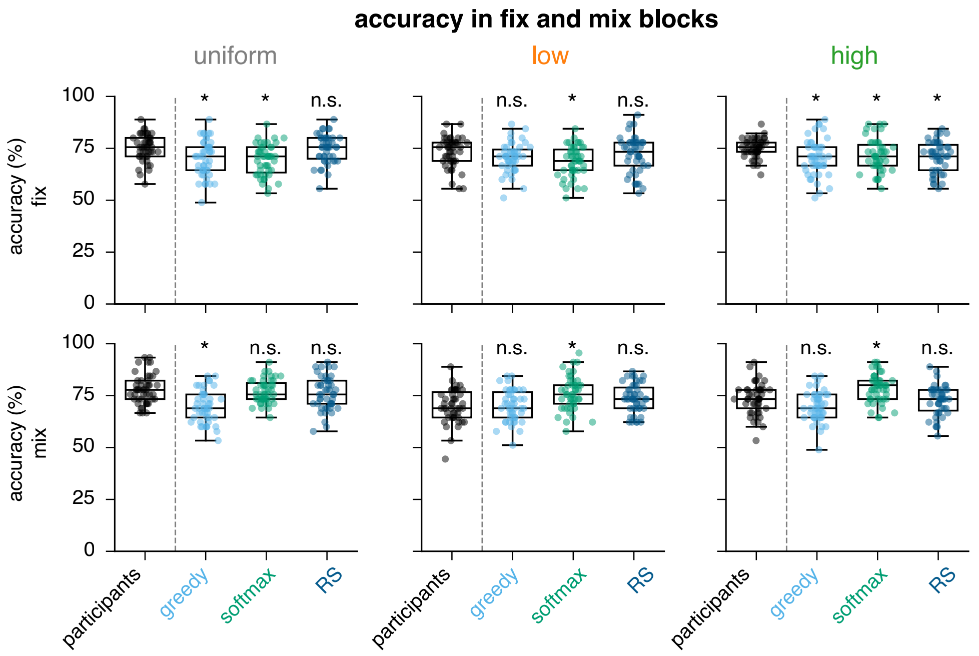


**Fig. S4** Accuracy (%) for uniform, low, and high decks in fix (top row) and mix (bottom row) blocks for participants and each model. Participants vs. model comparisons are displayed above each model panel (* $p < 0.05$, n.s: not significant)

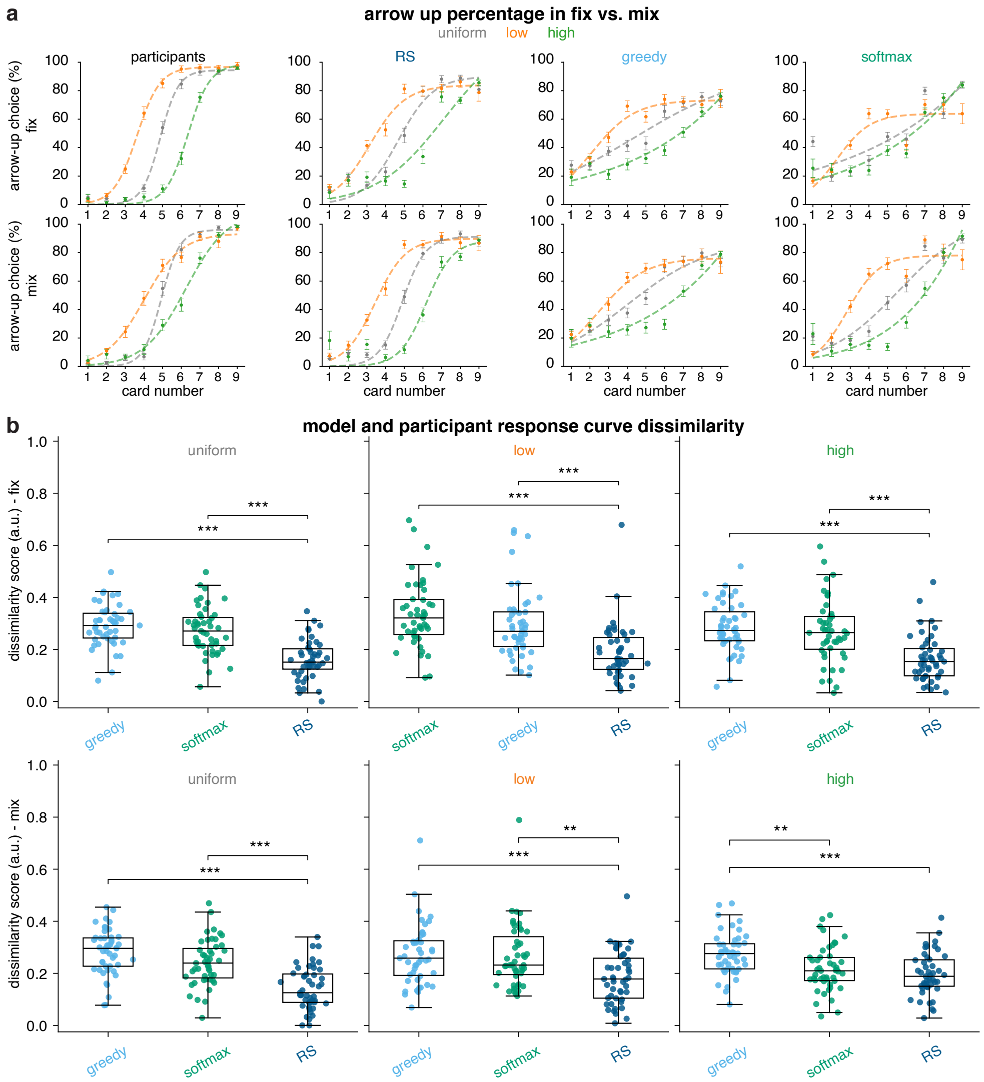


**Fig. S5** (**a**) Arrow-up choice percentages across deck and card number in fix (top row) and mix (bottom row) blocks, comparing participant and model choices. (**b**) Response-curve dissimilarity scores between participants and models for each card decks (uniform, low, high) in fix (top row) and mix (bottom row) blocks (* $p < 0.05$, ** $p < 0.01$, *** $p < 0.001$).

**Table S1** Risk for each *card*1 (participant's card) in the uniform deck.

| $card1$ | $N(card1)$ | $45 - N(card1)$ | $p(card2 < card1)$ | $p(card2 > card1)$ | Risk |
|---|---|---|---|---|---|
| 1 | 5 | 40 | 0.000 | 1.000 | 0.000 |
| 2 | 5 | 40 | 0.125 | 0.875 | 0.125 |
| 3 | 5 | 40 | 0.250 | 0.750 | 0.250 |
| 4 | 5 | 40 | 0.375 | 0.625 | 0.375 |
| 5 | 5 | 40 | 0.500 | 0.500 | 0.500 |
| 6 | 5 | 40 | 0.625 | 0.375 | 0.375 |
| 7 | 5 | 40 | 0.750 | 0.250 | 0.250 |
| 8 | 5 | 40 | 0.875 | 0.125 | 0.125 |
| 9 | 5 | 40 | 1.000 | 0.000 | 0.000 |

**Table S2** Risk for each *card*1 (participant's card) in the low deck.

| $card1$ | $N(card1)$ | $45 - N(card1)$ | $p(card2 < card1)$ | $p(card2 > card1)$ | Risk |
|---|---|---|---|---|---|
| 1 | 9 | 36 | 0.000 | 1.000 | 0.000 |
| 2 | 8 | 37 | 0.243 | 0.757 | 0.243 |
| 3 | 7 | 38 | 0.447 | 0.553 | 0.447 |
| 4 | 6 | 39 | 0.615 | 0.385 | 0.385 |
| 5 | 5 | 40 | 0.750 | 0.250 | 0.250 |
| 6 | 4 | 41 | 0.854 | 0.146 | 0.146 |
| 7 | 3 | 42 | 0.929 | 0.071 | 0.071 |
| 8 | 2 | 43 | 0.977 | 0.023 | 0.023 |
| 9 | 1 | 44 | 1.000 | 0.000 | 0.000 |

**Table S3** Risk for each *card*1 (participant's card) in the high deck.

| $card1$ | $N(card1)$ | $45 - N(card1)$ | $p(card2 < card1)$ | $p(card2 > card1)$ | Risk |
|---|---|---|---|---|---|
| 1 | 1 | 44 | 0.000 | 1.000 | 0.000 |
| 2 | 2 | 43 | 0.023 | 0.977 | 0.023 |
| 3 | 3 | 42 | 0.071 | 0.929 | 0.071 |
| 4 | 4 | 41 | 0.146 | 0.854 | 0.146 |
| 5 | 5 | 40 | 0.250 | 0.750 | 0.250 |
| 6 | 6 | 39 | 0.385 | 0.615 | 0.385 |
| 7 | 7 | 38 | 0.447 | 0.553 | 0.447 |
| 8 | 8 | 37 | 0.243 | 0.757 | 0.243 |
| 9 | 9 | 36 | 0.000 | 1.000 | 0.000 |

**Table S4** Linear regression results for choice across RS, Greedy, and Softmax models. Rows with $p < 0.05$ are in bold.

| model | predictor | coef | std-err (clustered) | z | p |
|---|---|---|---|---|---|
| RS | baseline | 0.294855 | 0.120822 | 2.4402 | 0.0880 |
| | $\|\Delta Q\|$ | **-0.091563** | **0.010892** | **-8.4026** | **<0.001** |
| | **deck - high** | **0.241338** | **0.037410** | **6.4512** | **<0.001** |
| | **deck - low** | **0.273379** | **0.037635** | **7.2639** | **<0.001** |
| | $\|5 - \textbf{card}\|$ | **-0.255921** | **0.031071** | **-8.2376** | **<0.001** |
| | **log(trial)** | **-0.100699** | **0.031593** | **-3.1869** | **0.0010** |
| Greedy | baseline | 0.260615 | 0.200887 | 1.2967 | 0.1956 |
| | $\|\Delta Q\|$ | **-0.036174** | **0.013870** | **-2.6084** | **0.0091** |
| | **deck - high** | **0.267142** | **0.041681** | **6.4093** | **<0.001** |
| | **deck - low** | **0.288583** | **0.041373** | **6.9724** | **<0.001** |
| | $\|5 - \textbf{card}\|$ | **-0.262979** | **0.030742** | **-8.5540** | **<0.001** |
| | **log(trial)** | **-0.097196** | **0.030139** | **-3.2253** | **0.0012** |
| Softmax | baseline | 0.264129 | 0.192614 | 1.3707 | 0.1699 |
| | $\|\Delta Q\|$ | **-0.040192** | **0.013890** | **-2.8930** | **0.0038** |
| | **deck - high** | **0.272669** | **0.041491** | **6.5723** | **<0.001** |
| | **deck - low** | **0.288234** | **0.041419** | **6.9575** | **<0.001** |
| | $\|5 - \textbf{card}\|$ | **-0.262051** | **0.030838** | **-8.4978** | **<0.001** |
| | **log(trial)** | **-0.098116** | **0.030340** | **-3.2349** | **0.0012** |

**Table S5** Logistic regression results for choice across RS, Greedy, and Softmax models. Rows with $p < 0.05$ are in bold.

| model | predictor | coef | std-err (clustered) | z | p |
|---|---|---|---|---|---|
| RS | baseline | -0.181 | 0.145 | -1.246 | 0.211 |
| | $\Delta Q$ | **5.765** | **0.773** | **7.454** | $<$**0.001** |
| | **deck - high** | **0.597** | **0.085** | **7.023** | $<$**0.001** |
| | **deck - low** | **-0.280** | **0.091** | **-3.087** | **0.002** |
| | log(trial) | 0.033 | 0.034 | 0.970 | 0.333 |
| Greedy | baseline | -0.113 | 0.110 | -1.018 | 0.308 |
| | $\Delta Q$ | **4.824** | **0.667** | **7.230** | $<$**0.001** |
| | **deck - high** | **0.343** | **0.082** | **4.179** | $<$**0.001** |
| | **deck - low** | **-0.285** | **0.083** | **-3.422** | $<$**0.001** |
| | log(trial) | 0.014 | 0.028 | 0.506 | 0.612 |
| Softmax | baseline | -0.066 | 0.097 | -0.685 | 0.494 |
| | $\Delta Q$ | **4.432** | **0.696** | **6.368** | $<$**0.001** |
| | deck - high | 0.159 | 0.124 | 1.266 | 0.205 |
| | **deck - low** | **-0.257** | **0.076** | **-3.374** | $<$**0.001** |
| | log(trial) | 0.038 | 0.031 | 1.259 | 0.208 |

**Table S6** Demographic characteristics of epileptic and healthy participants. Age is reported in categorical bins; gender reflects self-reported categories.

| Category | Level | Epileptic (N = 10) | Healthy (N = 37) |
|---|---|---|---|
| Age | 18–24 | 2 | 5 |
| | 25–34 | 4 | 27 |
| | 35–44 | 3 | 1 |
| | 45–54 | 1 | 2 |
| | 65+ | 0 | 2 |
| Gender | Female | 4 | 18 |
| | Male | 6 | 16 |
| | Non-binary | 0 | 2 |
| | Prefer not to say | 0 | 1 |
